# DNA Features Viewer, a sequence annotations formatting and plotting library for Python

**DOI:** 10.1101/2020.01.09.900589

**Authors:** Valentin Zulkower, Susan Rosser

## Abstract

**Motivation:** While the Python programming language counts many Bioinformatics and Computational Biology libraries, none offers customizable sequence annotations visualizations with layout optimization.

**Results:** DNA Features Viewer is a sequence annotations plotting library which optimizes plot readability while letting users tailor other visual aspects (colors, labels, highlights, etc.) to their particular use case.

**Availability:** Open-source code and documentation are available on Github under the MIT licence (https://github.com/Edinburgh-Genome-Foundry/DnaFeaturesViewer).

**Supplementary information:** attached.

## 1 Introduction

DNA sequence visualization is a common need in Bioinformatics, and many software tools can plot sequence annotations from Genbank or General Feature Format (GFF) records. A sequence annotation specifies a location (start position, end position and strand), feature type (such as “CDS” or “regulatory”) and attributes (e.g. gene name, species of origin, or locus tag). When displaying a record with many annotations, one may want to enhance readability by hiding or highlighting certain features and attributes to focus the reader’s attention on the most relevant information.

Interactive sequence editing software such as SnapGene Viewer (www.snapgene.com) or Benchling (www.benchling.com) enable users to manually color or hide sequence features, but the customization is limited and cannot be automated. Python modules for sequence plotting are scarce and lack automation capabilities, making them difficult to integrate with other projects (see Supplementary Section A for a review). For instance, both DnaPlotLib (Der *et al*., 2017) and Biopython (Cock *et al*., 2009) require users to style each annotation separately, and do not automatically avoid collisions between overlapping annotations and their labels.

Here we present DNA Features Viewer, a Python library which lets users define visual “themes” determining the label and display style of each annotation as a function of its type, location, and attributes. Annotations are then automatically laid out to create compact and readable plots, making the library a robust choice as a generic plotter for other frameworks. Plots can be exported in PNG, SVG, PDF or interactive HTML format, for use in interactive notebooks, PDF reports, or web applications.

## 2 Usage and examples

### 2.1 Definition of visual themes

In DNA Features Viewer, sequence annotation records read from Genbank or GFF files are converted to so-called *graphic records*, which define the visual aspects of each annotation. The conversion is ensured by a user-defined Python class (the *translator*) whose attributes and methods indicate which annotations should appear in the plot (and which should be discarded), as well as the visual style of each annotation, including arrow color, arrow width, edge width, label text, associated label in the figure’s legend, and text font properties. For instance, the translator class used in Figure 1A sets the label text as either the `\note` or `\gene` attribute of the annotation, assigns each feature’s color based on the feature’s type, and reports the color/type correspondence in the figure legend. A translator thus acts as a visual theme which can be defined once and used throughout a project to ensure style consistency across annotation plots.

**Fig. 1.**
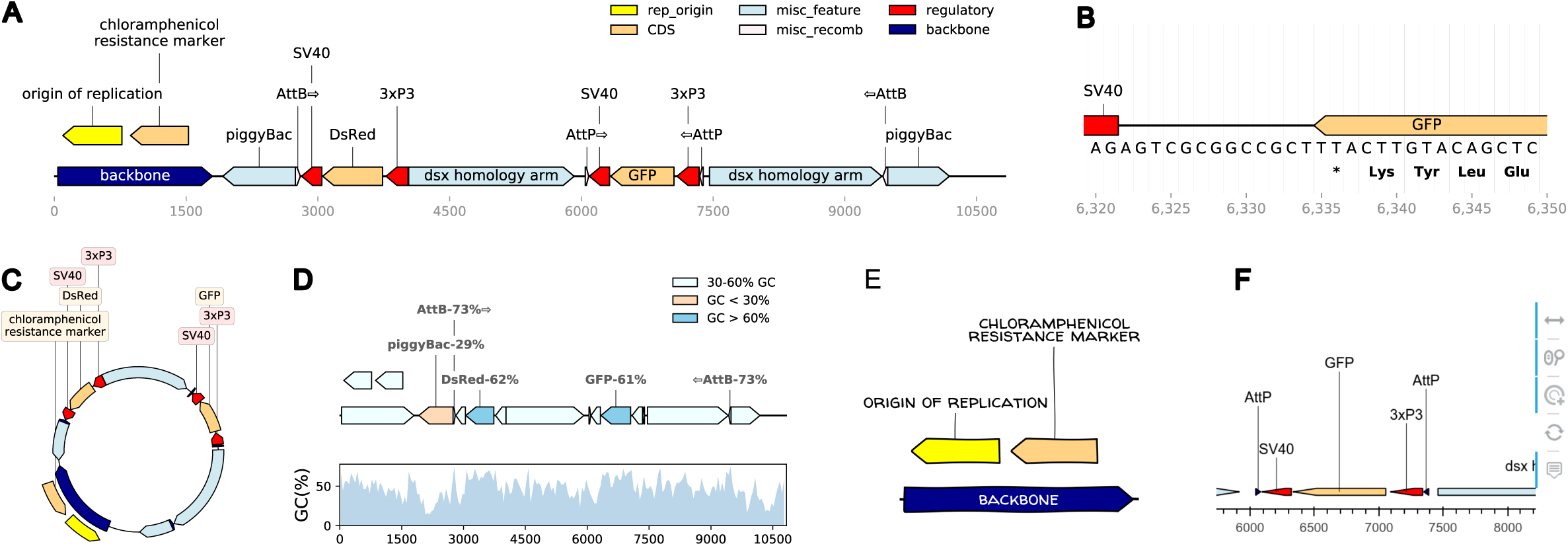
Different views of a pBac cloning vector (Kyrou et al., 2018) plotted using DNA Features viewer. The Python code to generate each figure is provided in Supplementary Section C. (A) Plasmid map plot using a custom visual theme as described in Section 2.1. (B) Detail plot focusing on a short sequence segment, with nucleotide and amino-acid sequences, and vertical visual guides. (C) Circular view of the plasmid. In this visual theme, label text boxes are automatically colored to be easily associated with their corresponding features. (D) Plasmid plot with colors indicating the GC content at each feature’s location. High- and low-GC features are highlighted with a label indicating their average GC content. The bottom subplot, which shares the same x-axis, indicates the local GC content over 100-nucleotides windows. (E) Plot using Matplotlib’s path.sketch filter and a custom font to create a “handwriting” effect. (F) Interactive HTML plot generated via the Bokeh library (shown here with a zoom around the position at location 7000). Icons on the left refer to widgets enabling mouse-based interactions.

### 2.2 Plot readability optimizations

Figure 1A also illustrates how DNA Feature Viewer automatically lays out the visual elements of a graphic record to optimize compactness and readability. Feature labels such as “backbone” and “GFP” are displayed directly inside their corresponding feature arrow, and the font color is automatically selected (as black or white) to fit the feature’s background color. Labels which do not fit inside a feature arrow are displayed above it, and wrapped on several lines when necessary (e.g. “chloramphenicol resistance marker”). For narrow features whose orientation cannot be easily discerned (such as AttB and AttP sites in Figure 1A), an arrow is added in the label. Finally, all features and label texts are organized along different vertical levels to avoid collisions (the layout optimization method, which uses variant of graph coloring algorithm, is described in Supplementary Section B). This ensures that the resulting plot remains readable irrespective of the figure’s width, which is set by the user and often constrained by space limitations on a web page or PDF report.

### 2.3 Other visualization formats

DNA Features Viewer supports a variety of plotting formats to suit different use cases. For instance, it enables to focus on on a small sequence region, displaying the nucleotide and amino-acid sequences (Figure 1B), or to plot the record’s full sequence over multi-line, multipage PDF documents (as shown in Supplementary Section D). A record can also be displayed with a circular topology, with text labels on the top (Figure 1C).

The library relies primarily on the Matplotlib plotting framework (Hunter, 2007) for graphics rendering, making it possible to display sequence annotations along with other other data visualization. For instance DNA Features Viewer has been used to associate sequence maps with local ChIP RZ scores in Kroner *et al*. (2019), and local GC content in Greig *et al*. (2018) (also illustrated in Figure 1D). Matplotlib also allows to finely tune plotting style with custom fonts and path filters, as illustrated in Figure 1E, to suit different media (articles, presentation slides, etc.)

Finally, the Bokeh library (Bokeh Development Team, 2019) can be used as a plotting backend, although this support is limited to linear sequence views. This allows the rendition of graphic records as interactive HTML plots which can be integrated in a webpage and allow the exploration of very large features record thanks to interactive widgets to pan and zoom around local regions (as shown in Figure 1F).

## 3 Implementation

DNA Features Viewer is written in Python. Genbank file parsing is provided by the Biopython library, and GFF parsing by the BCBB library (https://github.com/chapmanb/bcbb, unpublished).

## Acknowledgments

We thank Yu-jin Kim for comments and suggestions.

## Supplementary Information to

### A. Other annotation plotting frameworks

In this section we compare different Python sequence annotation plotting frameworks to DNA Features Viewer. As a benchmark we use a GFF annotations file featuring 3 gene expression units, as shown in Table SI1, and we will show how each framework plots the record with minimal configuration.

**Table SI1:**
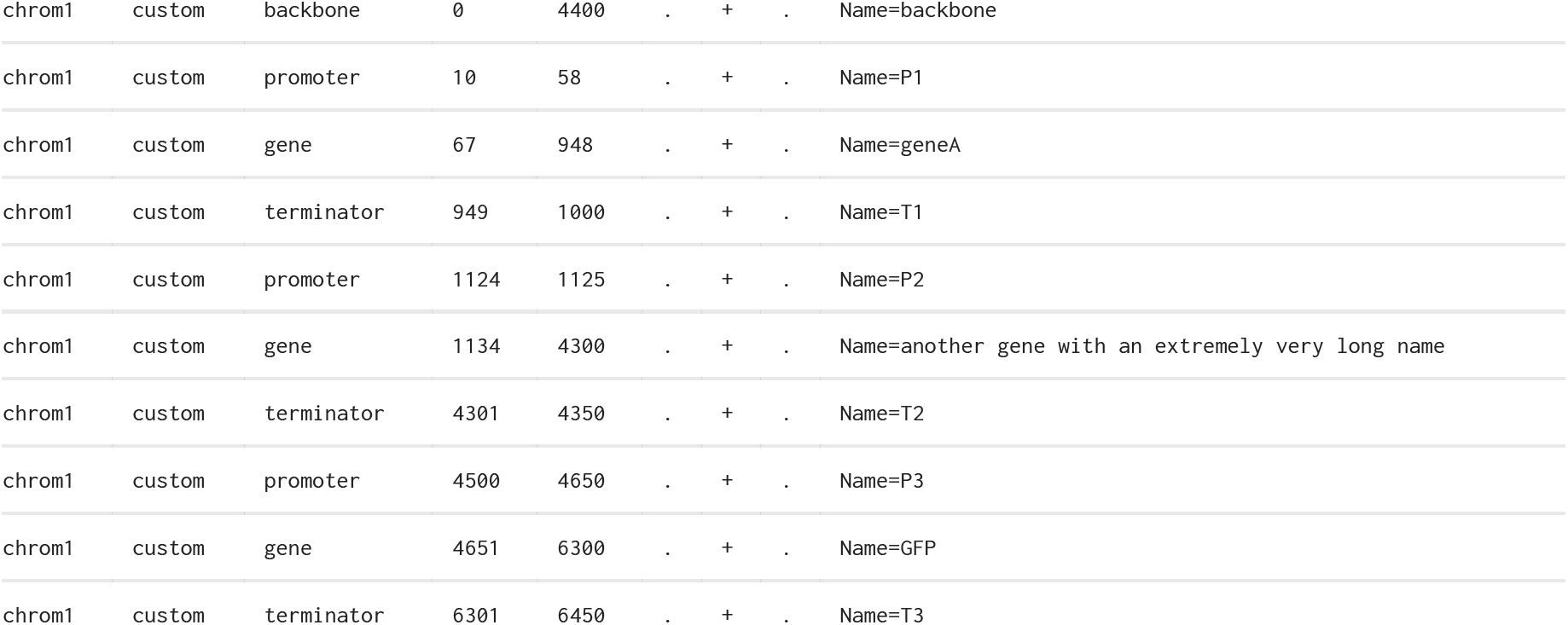
Annotations in the `plasmid.gff` file used as a benchmark in this section (the actual file contains exactly this information, with one entry per line and tabulations separating each entry’s columns).

|  |  |  |  |  |  |  |  |  |
| --- | --- | --- | --- | --- | --- | --- | --- | --- |
| chrom1 | custom | backbone | 0 | 4400 | . | + | . | Name=backbone |
| chrom1 | custom | promoter | 10 | 58 | . | + | . | Name=P1 |
| chrom1 | custom | gene | 67 | 948 | . | + | . | Name=geneA |
| chrom1 | custom | terminator | 949 | 1000 | . | + | . | Name=T1 |
| chrom1 | custom | promoter | 1124 | 1125 | . | + | . | Name=P2 |
| chrom1 | custom | gene | 1134 | 4300 | . | + | . | Name=another gene with an extremely very long name |
| chrom1 | custom | terminator | 4301 | 4350 | . | + | . | Name=T2 |
| chrom1 | custom | promoter | 4500 | 4650 | . | + | . | Name=P3 |
| chrom1 | custom | gene | 4651 | 6300 | . | + | . | Name=GFP |
| chrom1 | custom | terminator | 6301 | 6450 | . | + | . | Name=T3 |

#### A1. Plotting with DNA Features Viewer

We first plot the record using DNA Feature Viewer, without any configuration or customization:

##### Code

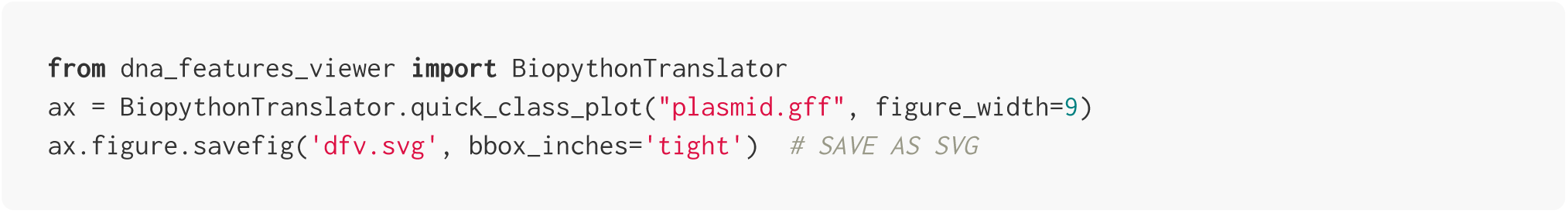

##### Result

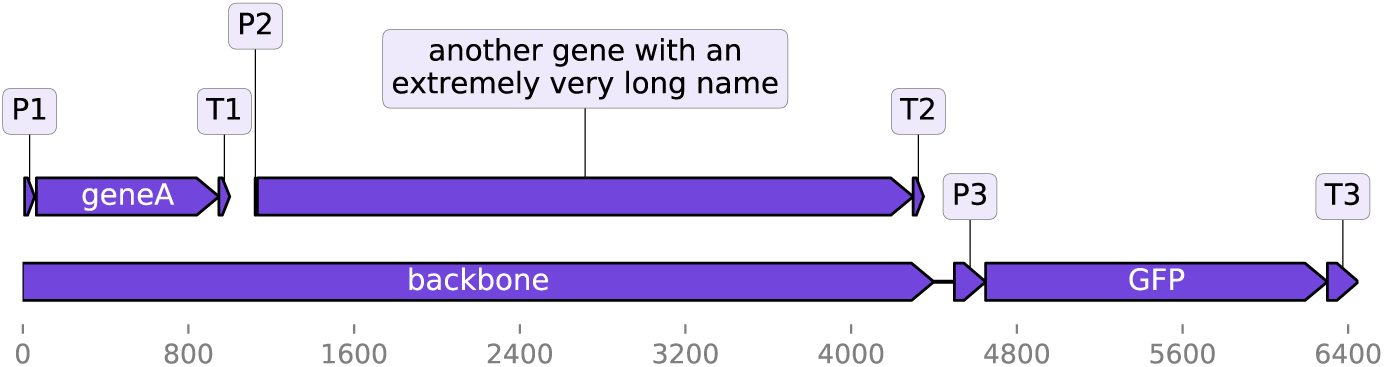

#### A2. Plotting with the Biopython plotting module

The script below is a variant from a script proposed in the official Biopython Cookbook tutorial (http://biopython.org/DIST/docs/tutorial/Tutorial.html):

##### Code

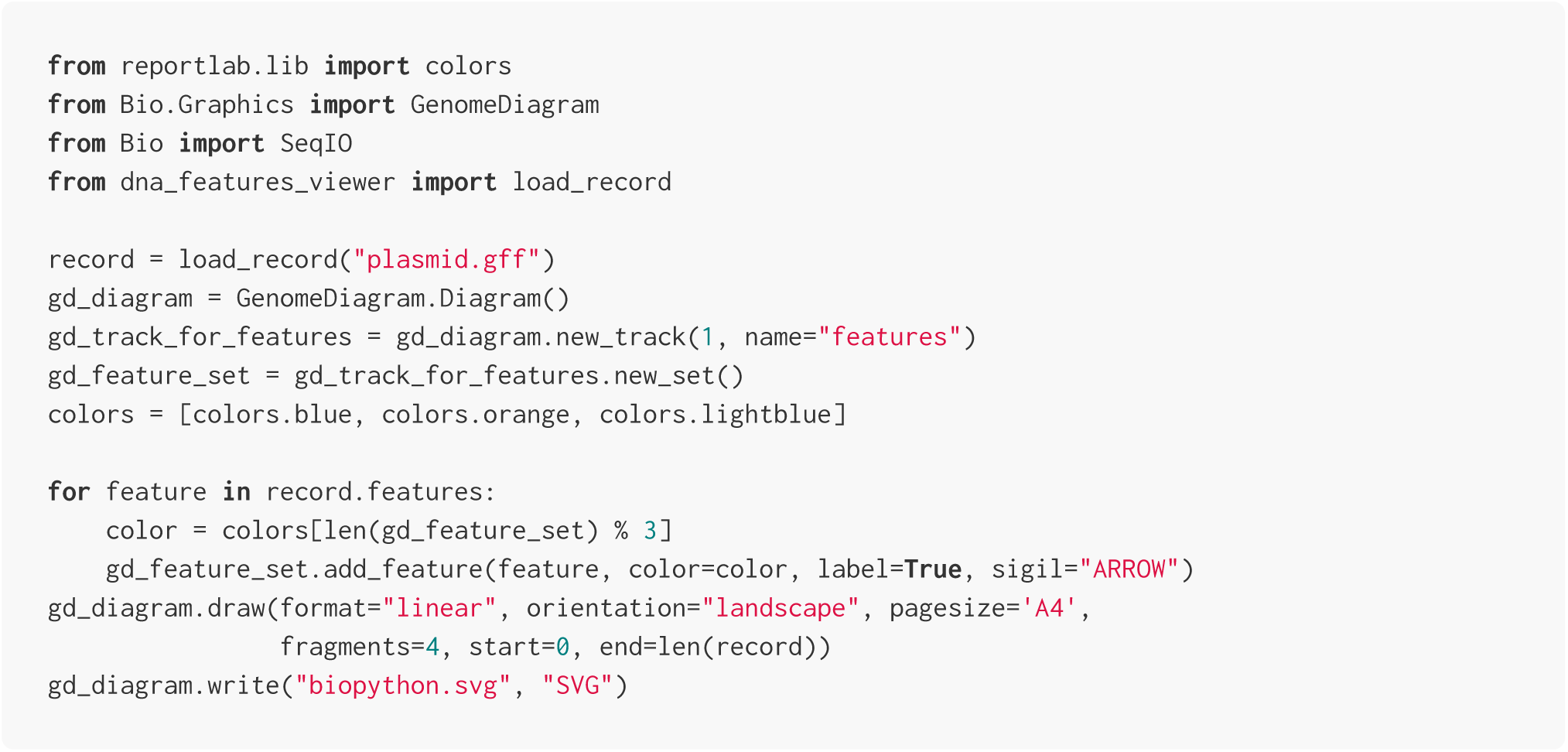

##### Result

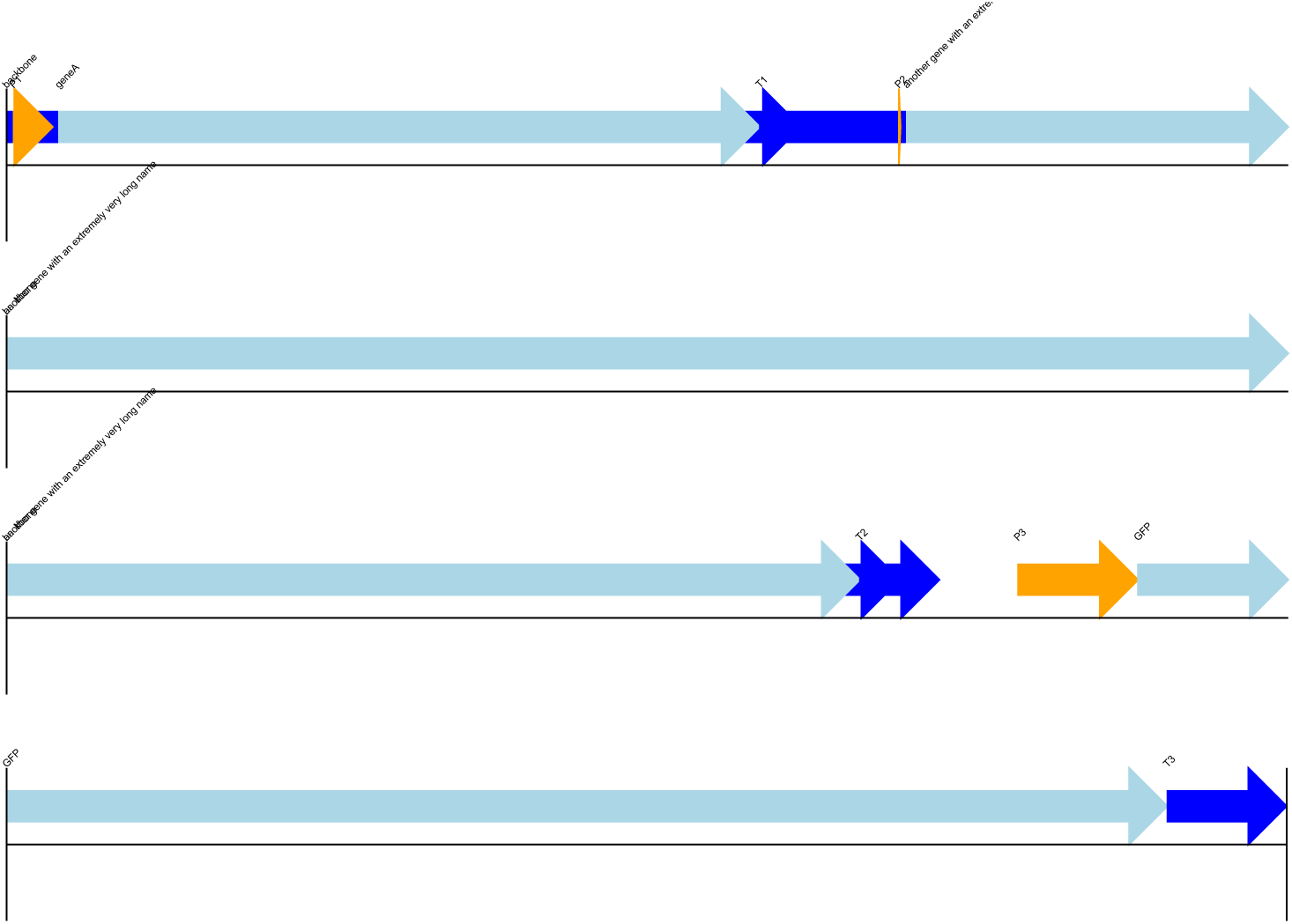

Biopython’s plotting module requires the user to specify colors for each feature separately (here we manually alternate between 3 colors, following the example of the Biopython tutorial, so that successive features can be distinguished). All features are plotted on a single line (unless the user places them manually on different tracks or different figures), causing overlapping features to collide. Labels are small, making the figure hard to read, although this also decreases the chances of label collisions (text collisions can still be seen at the beginning of lines 2 and 3).

#### A3. Plotting with DnaPlotLib

Here we use DnaPlotLib’s builtin `load_design_from_gff` method to plot the GFF file’s annotations with DnaPlotLib:

##### Code

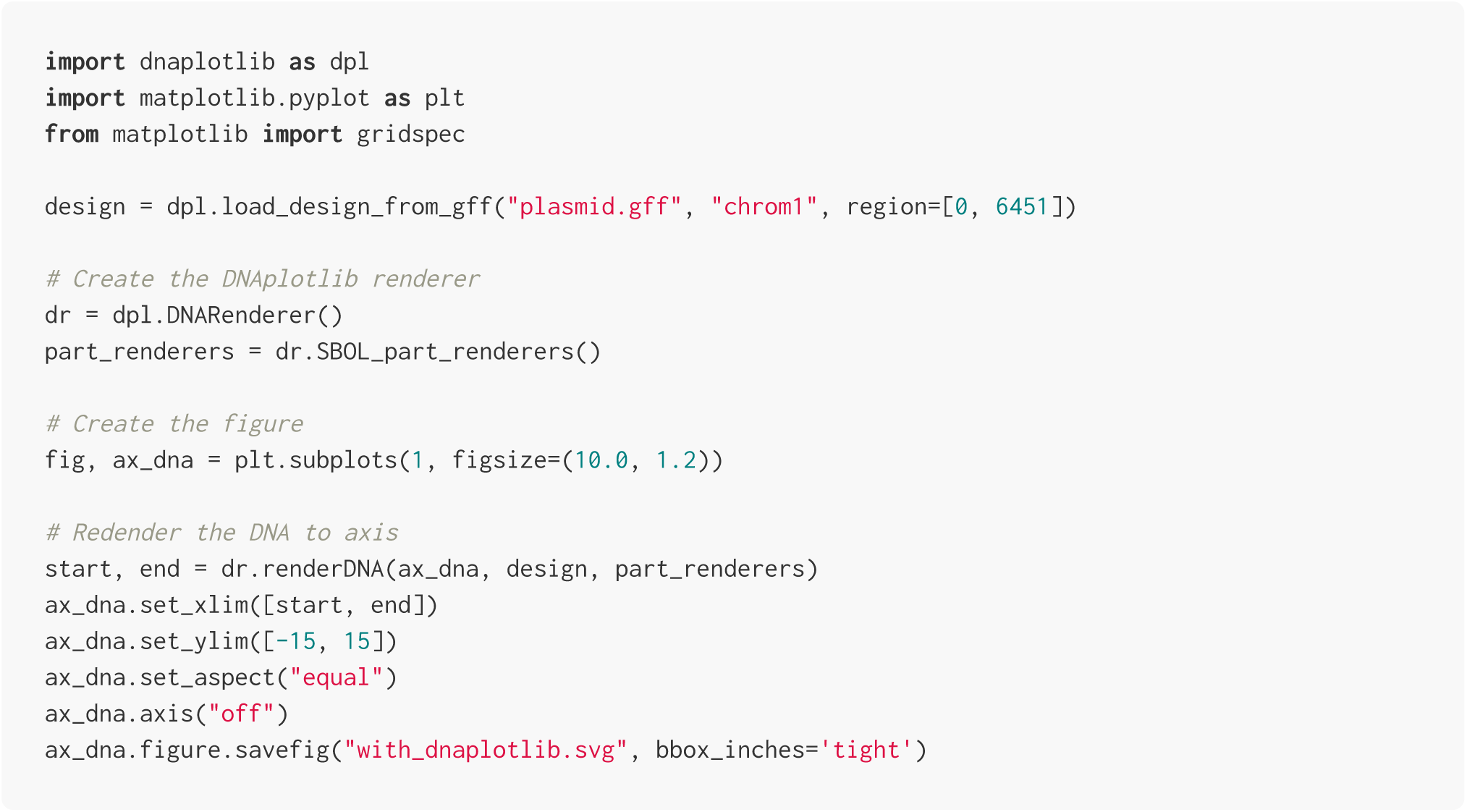

##### Result

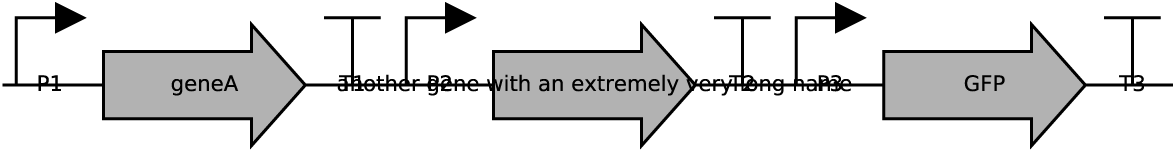

The DnaPlotLib library focuses on the display of the funtional genetic elements of a sequence (promoters, coding sequences, terminators, etc), and allows a high level of manual customization to produce publication-quality plots. However, we see in this example that it less adapted to the general display of annotations from arbitrary GFF or genbank records. The *“backbone”* annotation overlapping with the first two expression units is not displayed (information is lost) and collisions between text and other elements are not automatically avoided (some examples in the DnaPlotLib documentation show that it is possible to manually provide user-selected offsets in the Python script to place features and texts on different levels to avoid collisions, but this is not automated).

#### A4. The Biograpy library

The BiograPy library (A. Pierleoni, unpublished, http://apierleoni.github.io/BioGraPy/tutorial.html) and its most recent fork (M.O. Weber, unpublished, https://github.com/webermarcolivier/BioGraPy), allow users to define features which are then automatically placed in a plot so as to avoids collisions between overlapping features.

Unfortunately the libraries seem to rely on outdated dependencies (the latest code contributions to the projects are from 2016 and 2017) and we did not manage to run them on our example record. Therefore we are only showing screenshots from the projects’ websites in Figure SI1 below.

Among the notable differences with DNA Features Viewer, the labels are always placed inside or right under their corresponding feature’s arrow, which can be problematic for sequences with a high density of small annotations. The library does not feature any equivalent of DNA Features Viewer’s `BiopythonTranslator` to automatically convert genbank records to graphic records.

**Figure SI1:**
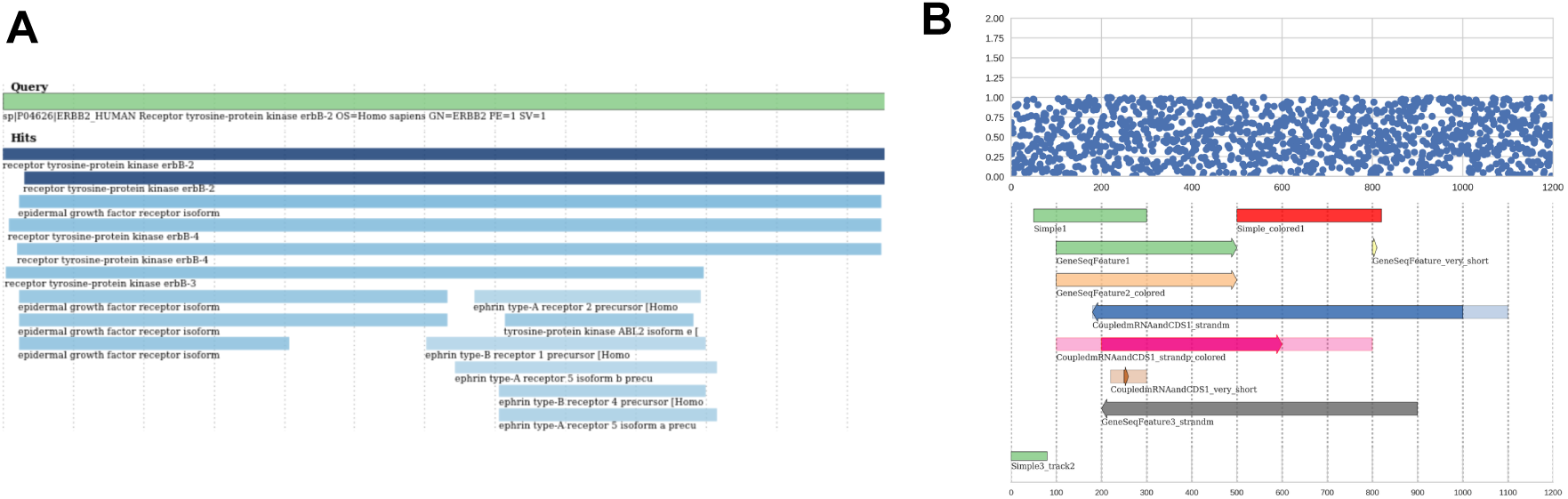
Sample outputs of the Biograpy library, generated by the original project (panel A) and its more recent fork (panel B). See the links provided in this paragraph for the source of these figures.

### B. Features and annotations positioning algorithm

#### Problem definition

This section describes how sequence features arrows and labels are vertically positioned by DNA Feature Viewer to avoid any collision (i.e. the superimposition of two graphical elements).

Every graphical element has a set horizontal coordinates (*x*_1_, *x*_2_). For feature arrows, these coordinates correspond to the feature’s start- and end-position in the sequence. For labels, it corresponds to the horizontal coordinates of the text after centering on the middle of the feature’s location. An element is said to be horizontally overlapping with another element at position 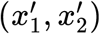 if the two segments overlap, which is equivalent to

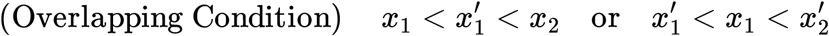

As of DNA Features Viewer v2.3, each element is placed vertically a on certain level *v* > 0, each level having approximately the same height as a line of text. Thus, two horizontally overlapping features placed will collide if and only if they are also placed on the same level.

The placement problem consists, for given a set of graphic elements, in determining a level *v* for each element, so that (1) no horizontally overlapping elements have the same level, and (2) the largest level *v* among all elements is as small as possible (to keep the plot compact).

#### Formulation as a graph coloring problem

The DNA Feature Viewer algorithm first builds a graph where each node represents a graphical element, and an edge between two nodes indicate that the corresponding elements are horizontally overlapping. The problem is now to find a *coloring* of each node with a level *v* that differs from the levels of all neighbors in the graph, to avoid collisions. This is a classical graph colorouring problem, which is known to be NP-complete (Brelax 1979), i.e. computationally intensive for large problems. Therefore, the algorithm uses *greedy coloring*, where it sequentially attributes the lowest available level to each element (starting with the widest elements so the larger features appear at the bottom of the plot):

- **For each** element:
  - List all the element’s neighbors in the graph for which a level has been set.
  - Set the element’s level to 0
  - **While** the element collides with any neighbor in the graph:
    ▪ Increase the element’s level by one.

Many small improvements are done to improve graph readability. First, larger features are considered before smaller ones in the iteration loop. As a result, the larger features always appear at the bottom of the plot. Second, all features arrows are attributed a level before all feature labels, so that the labels “float” on top of the feature arrows. Third, multi-line labels are taken into account, as explained in the next section.

#### Support for multi-line labels

Labels may have a certain number *N* of lines (feature arrows can be considered has being on a single line, i.e. *N* = 1). In this context, two horizontally overlapping elements, with respectively *N* lines at level *v*, and *N* ′ lines at *v*′, will collide if their levels are not sufficiently spread appart, or more precisely when the following condition (also illustrated in Figure SI2) is met:

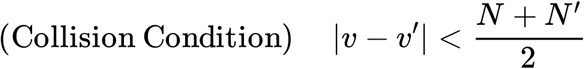

**Figure SI2:**
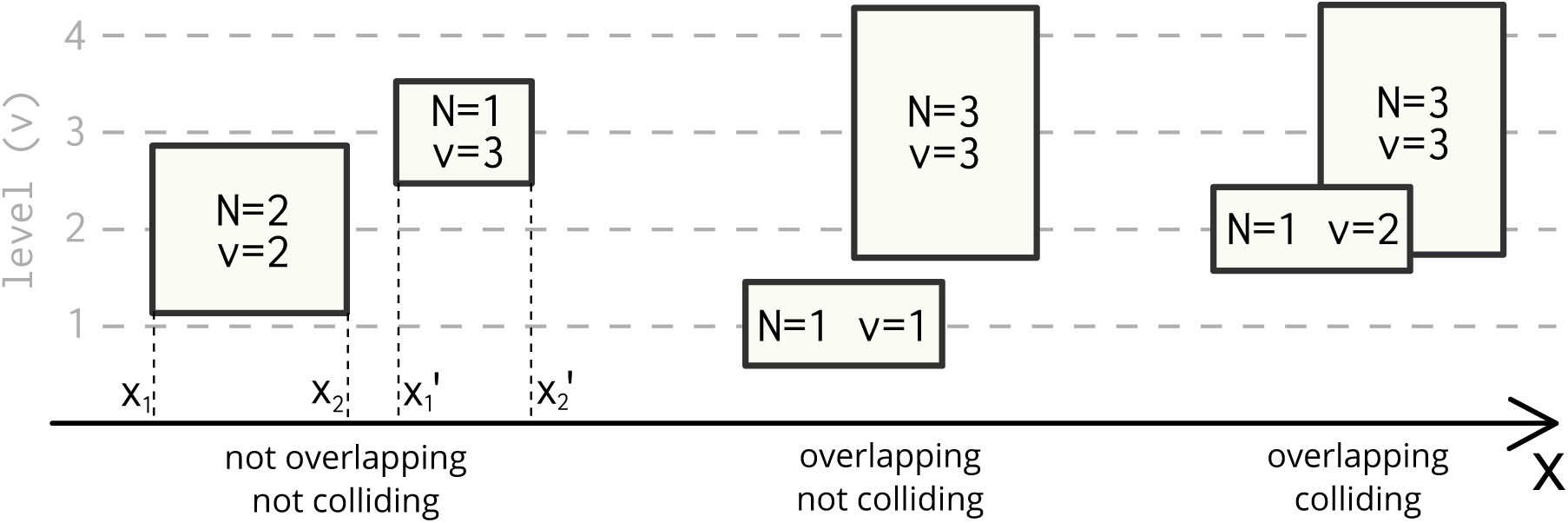
Graphical elements with varying level (v) and number of lines (N) as defined in the main text.

### C. Python code snippets for Figure 1

This sections provides the Python code for the plots shown in Figure 1 of the main text. Note that these snippets can also in the *Examples* section of the online project (https://github.com/Edinburgh-Genome-Foundry/DnaFeaturesViewer/tree/master/examples).

While the scripts to generate the different panels can be run in any order, they all require to first install the DNA Features Viewer library version 3.0 or more recent (either using the pip installation system via `pip install dna_features_viewer`, or by downloading and installing the library locally, as explained in the project documentation). It is also necessery to download (from NCBI.org) the plasmid record that will be used as a sample, using the code below:

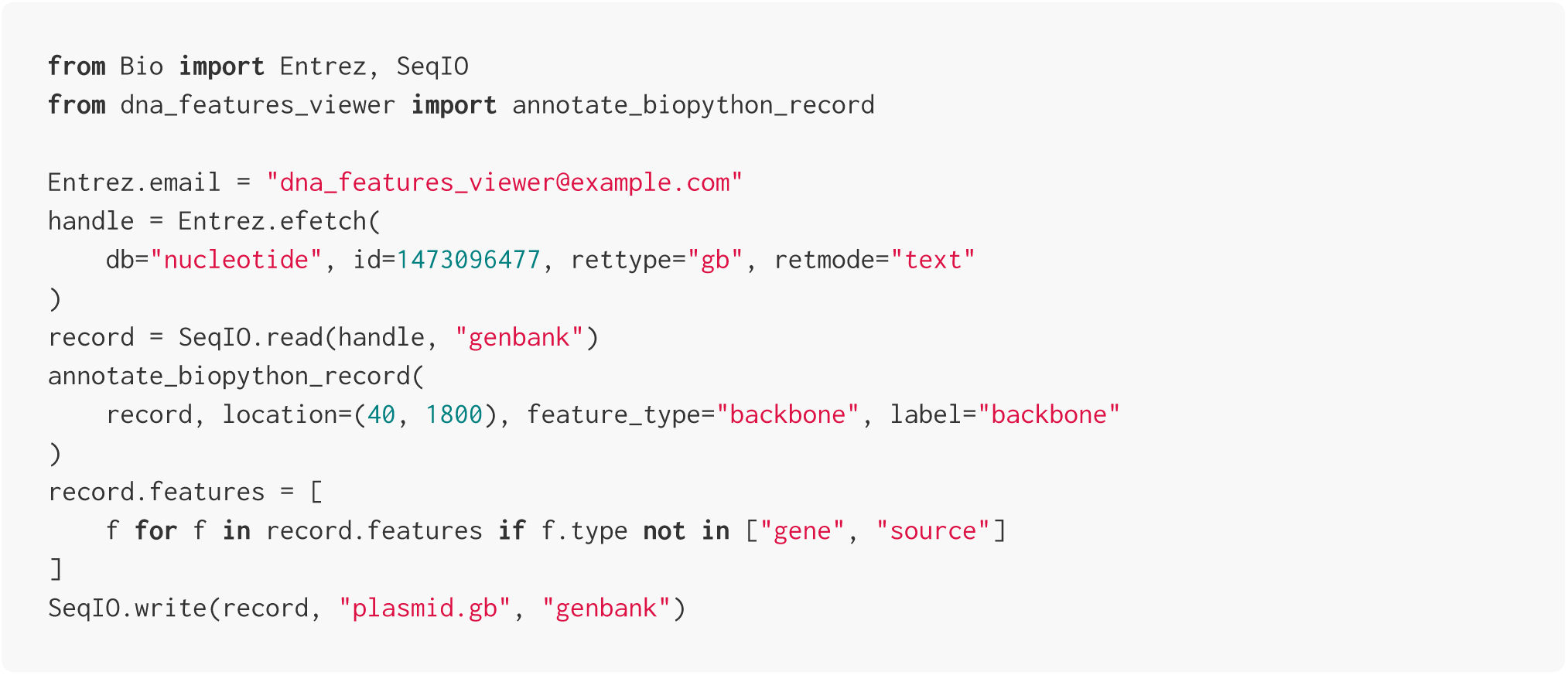

#### Panel A (linear view)

The script below defines a `CustomTranslator` class setting annotations colors based on feature type. All floating feature labels are written on a white background with no text box line.

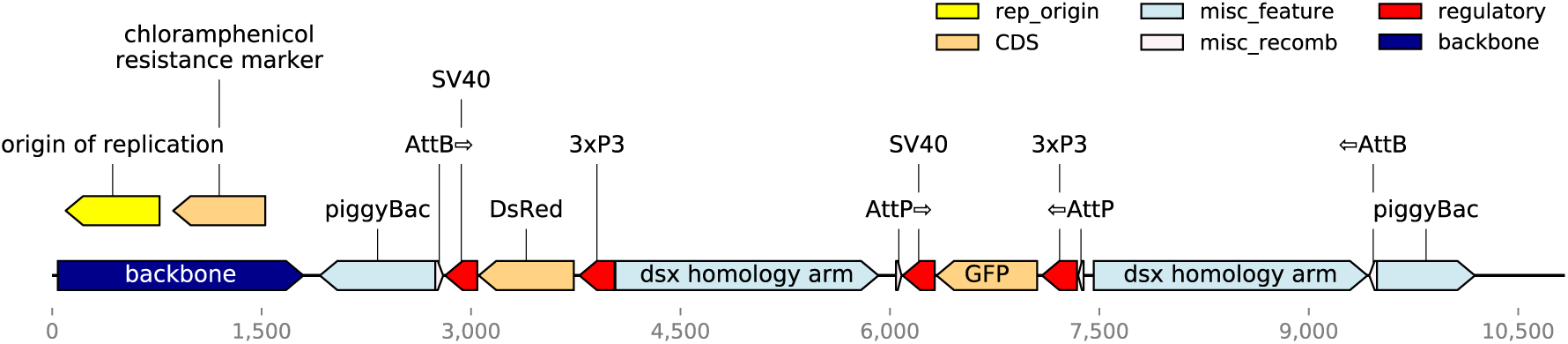

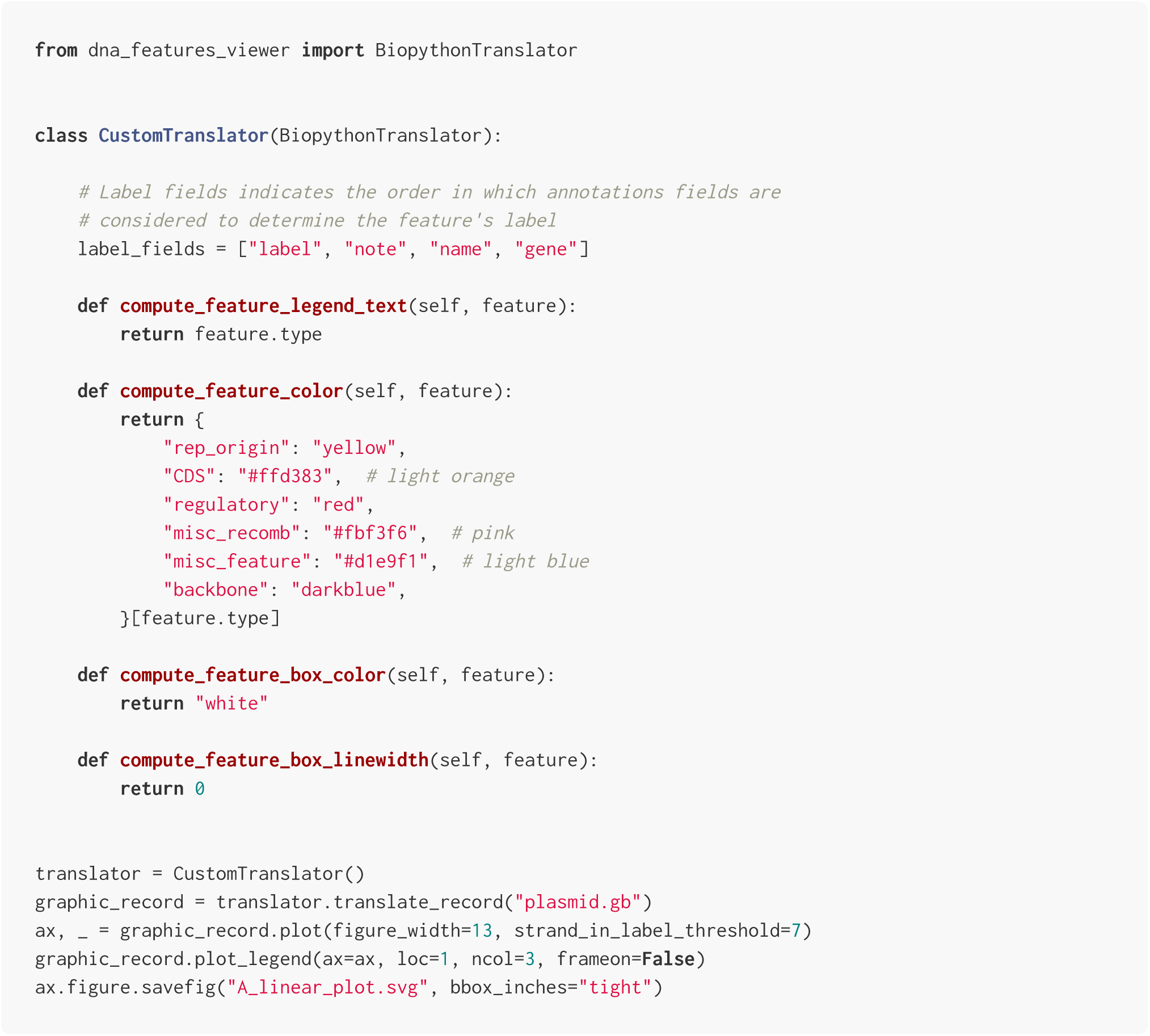

#### Panel B (detail view)

The script below imports the `CustomTranslator` defined in the previous section and uses it to display a cropped segment of the record (between indices 6320 and 6350), this time with nucleotide and amino-acid sequences overlaid on the figure.

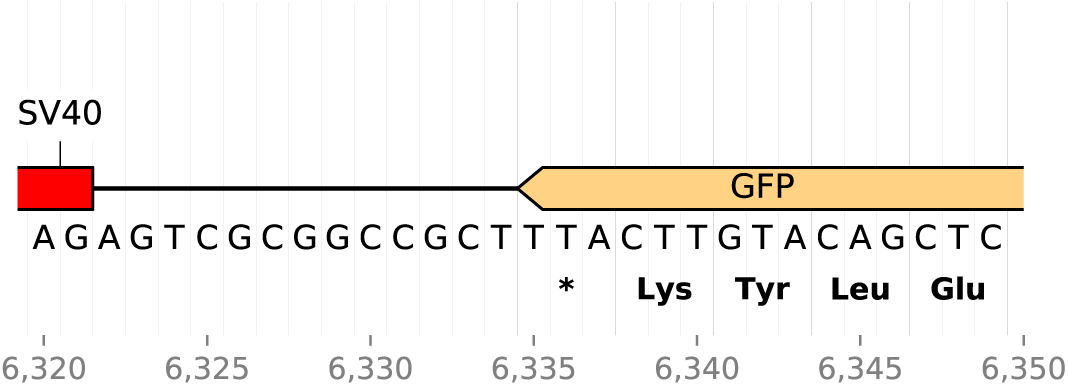

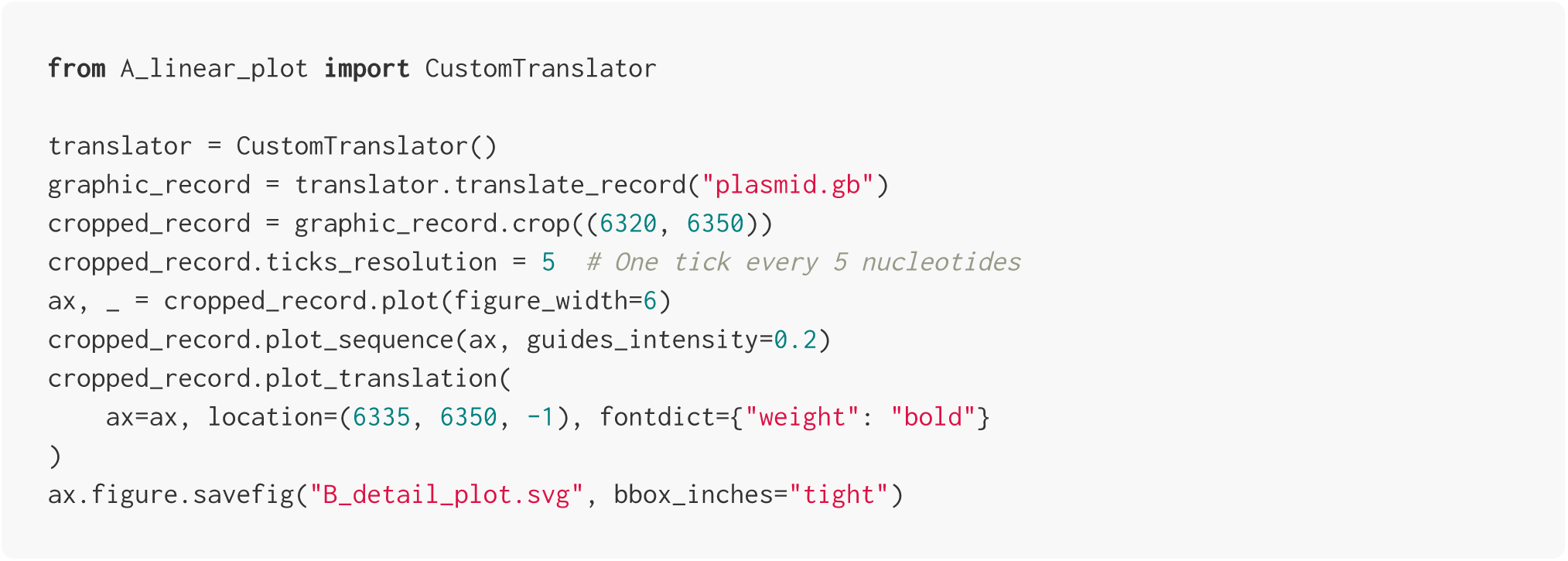

#### Panel C (circular view)

The script below defines an `ExpressionUnitTranslator` class which only labels coding sequences and regulatory elements. In addition, the script uses the `CircularGraphicRecord` class to plot the record, resulting in a circular plot.

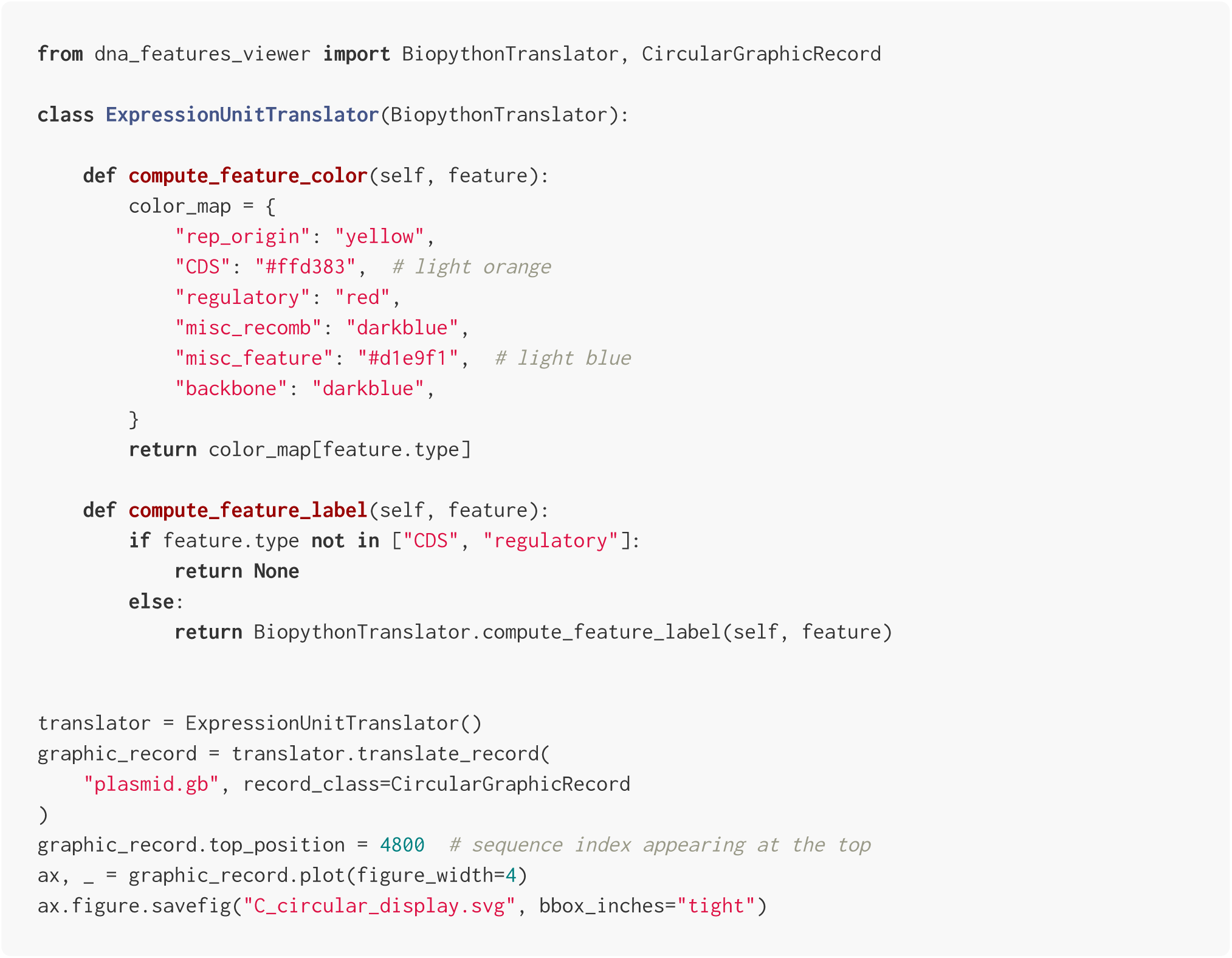

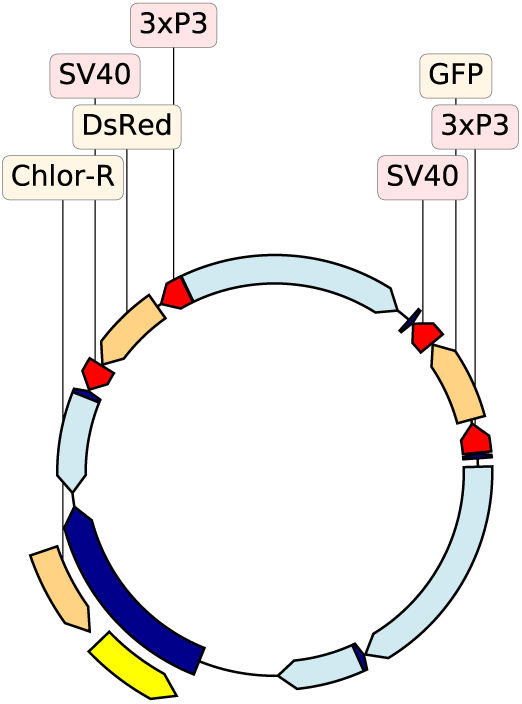

#### Panel D (GC% view)

The script below plots the sequence record alongside a profile of the local GC content. We first define a custom `GCIndicatingTranslator` class which separates features in regions with over 60% or under 30% GC, then we use Matplotlib to create two vertically-aligned subplots, for the record plot and GC content plot, respectively.

Note that `GCIndicatingTranslator` expects the features of the genbank record to be translated to have a `gc%` attribute. In the script, this attribute is computed for all features prior to creating the `GCIndicatingTranslator` instance.

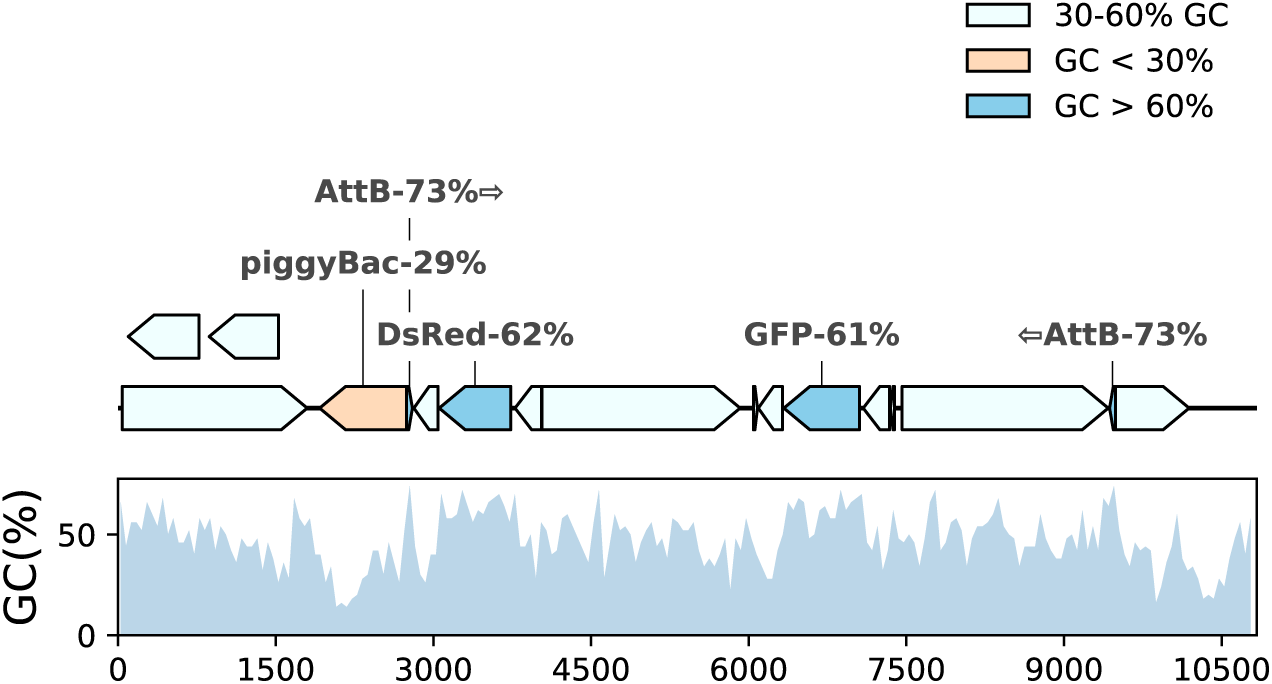

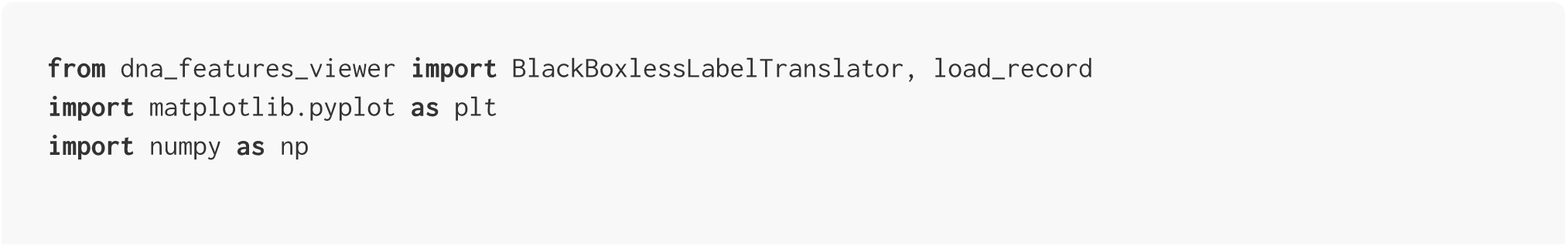

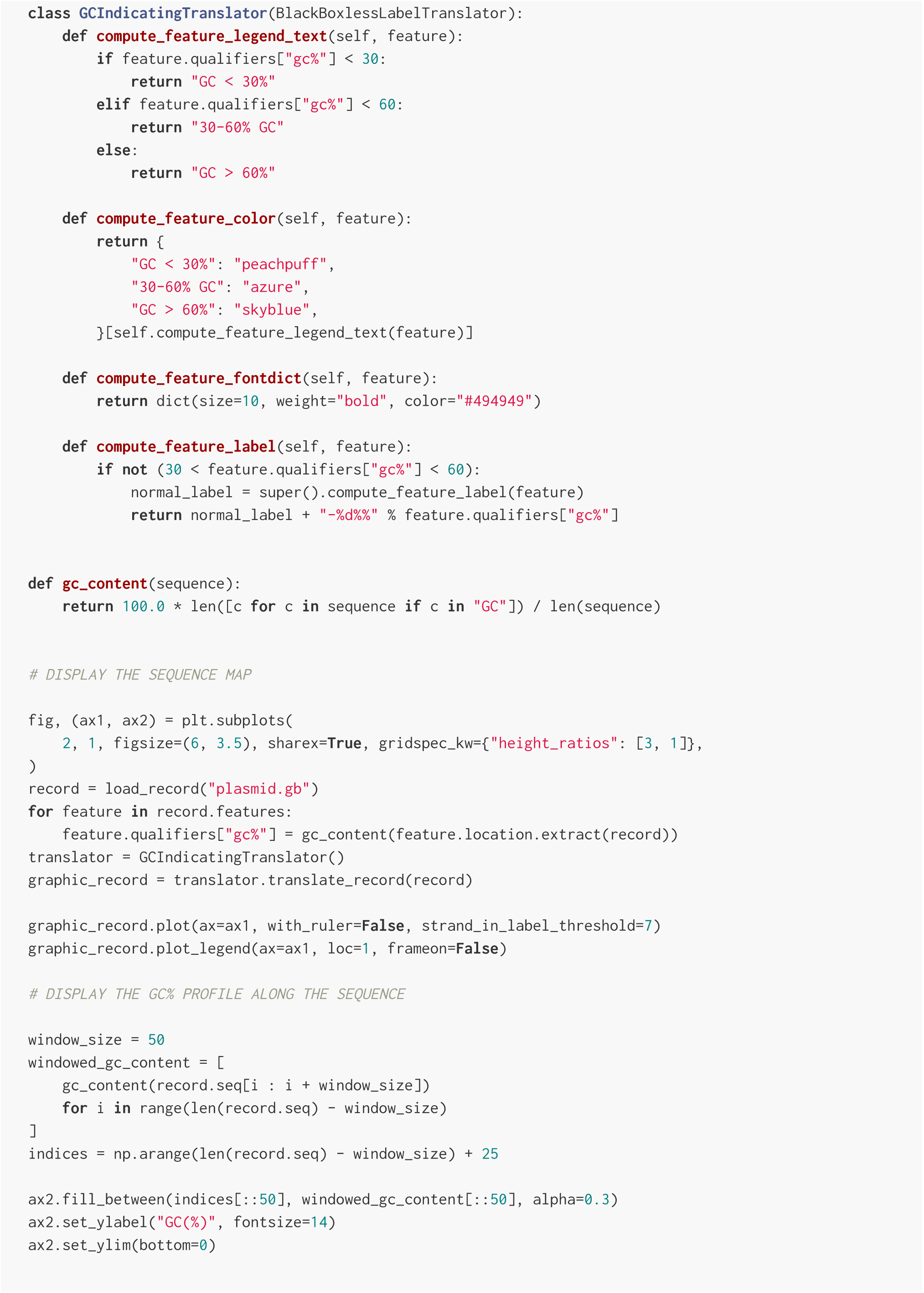

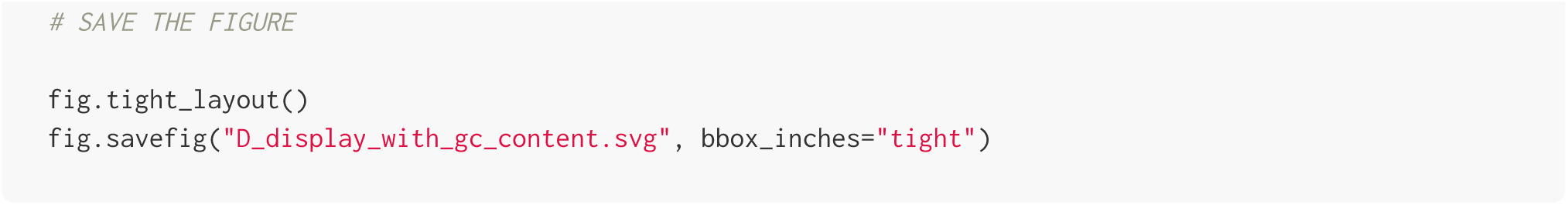

#### Panel E (sketch effect)

In this script we plot a cropped segment of the record, using a custom font, and the `path.sketch` filter of Matplotlib to introduce randomness in the line drawing, creating a *hand-drawing* effect.

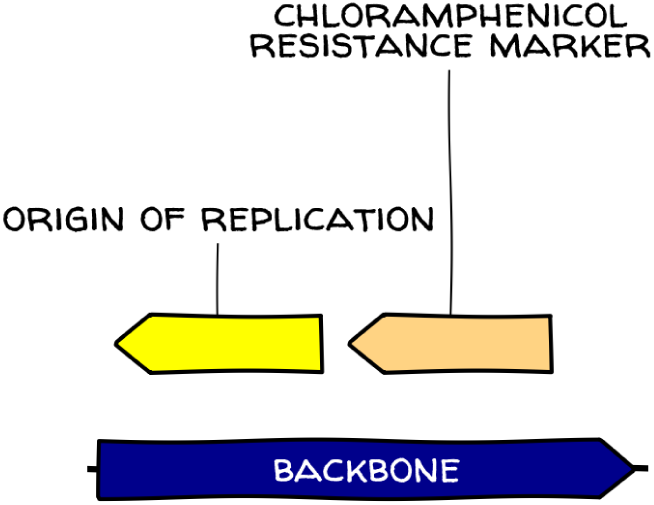

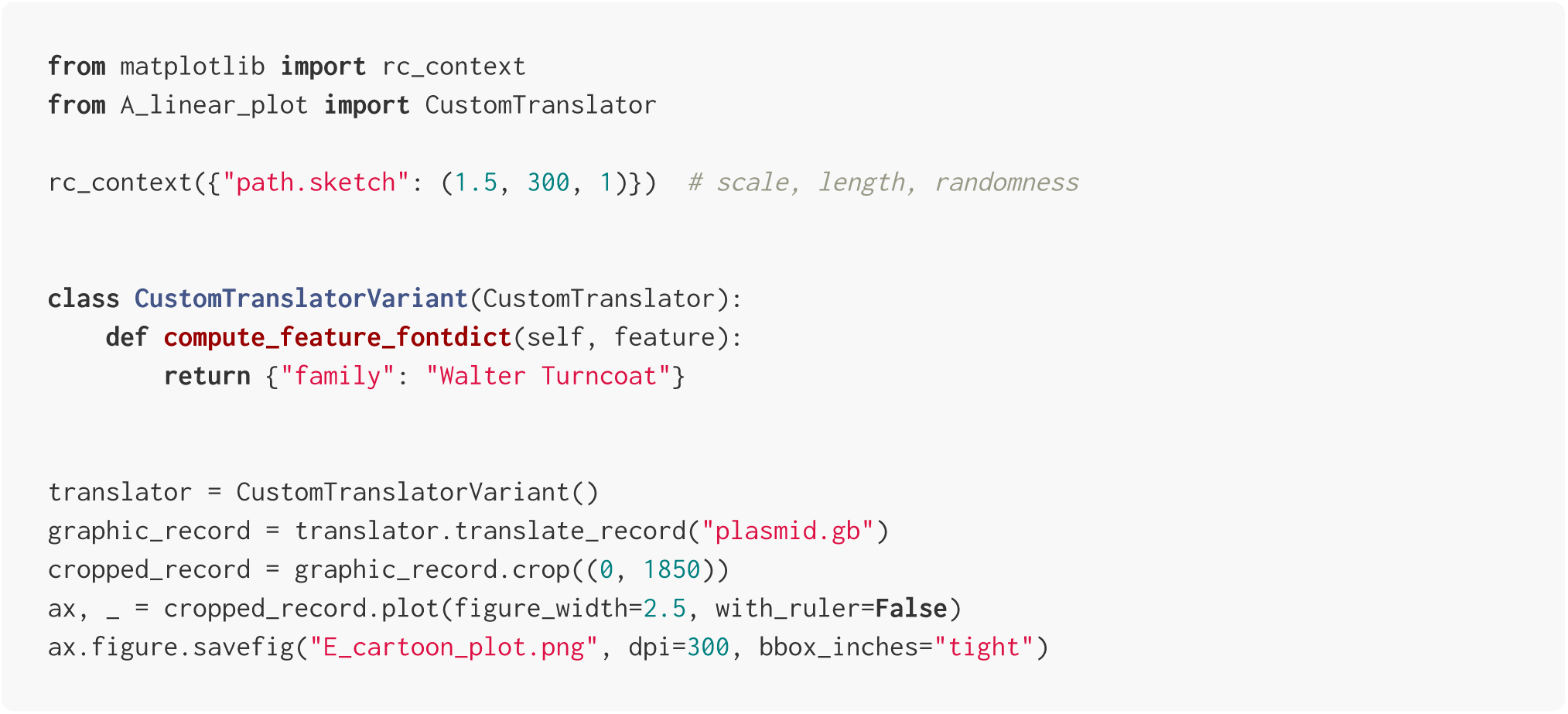

#### Panel F (interfactive plot)

In this script we create an HTML page featuring an interactive plot of the record using the Bokeh framework.

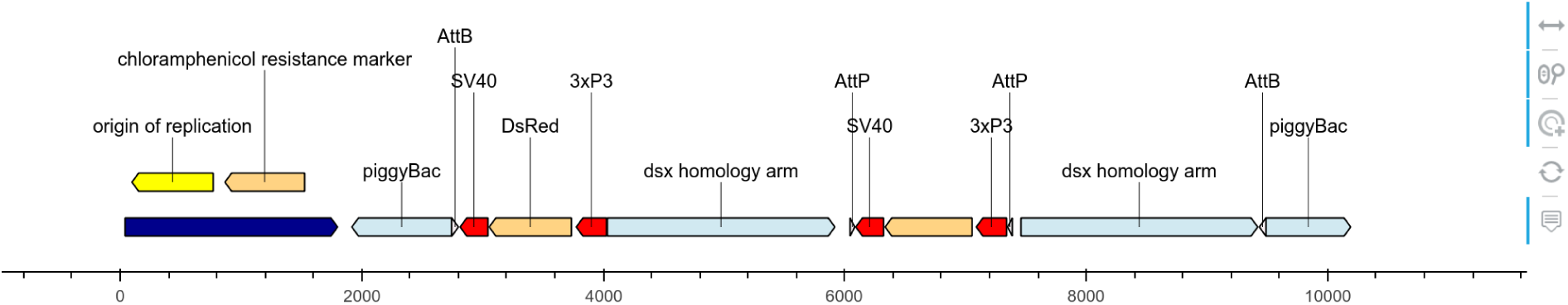

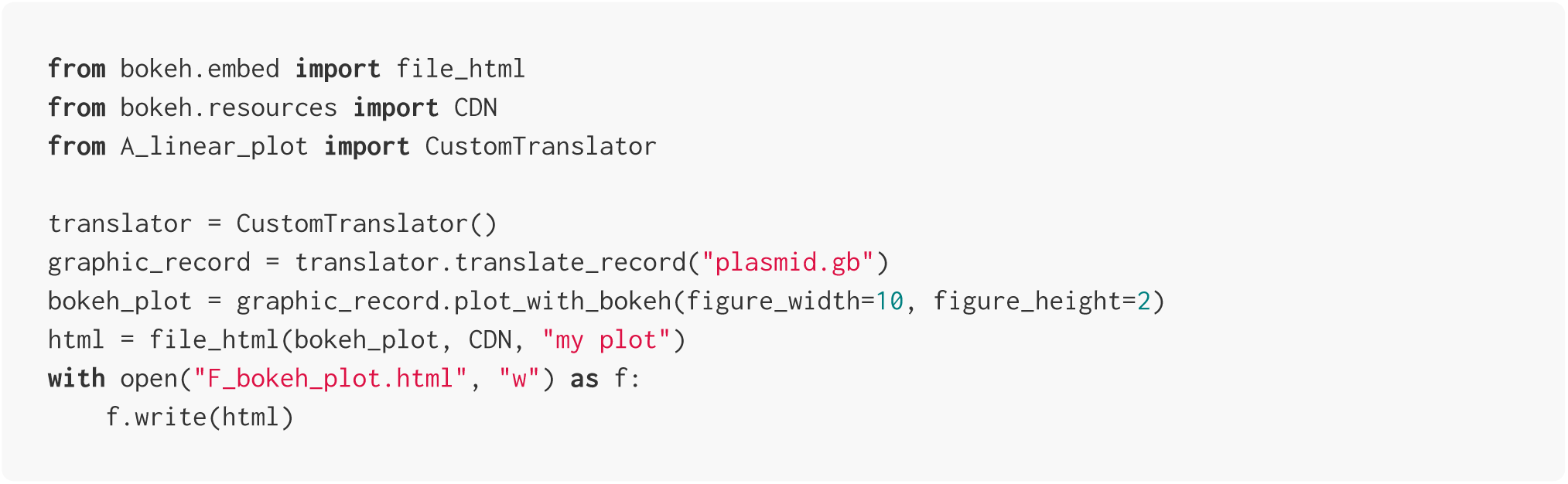

### D. Multiline, multi-page plot

The script below plots the full plasmid of the main text into a multi-line, multi-pages PDF file. The final file features 14 pages with 10 lines of 80 nucleotides per page (the three first pages are show below).

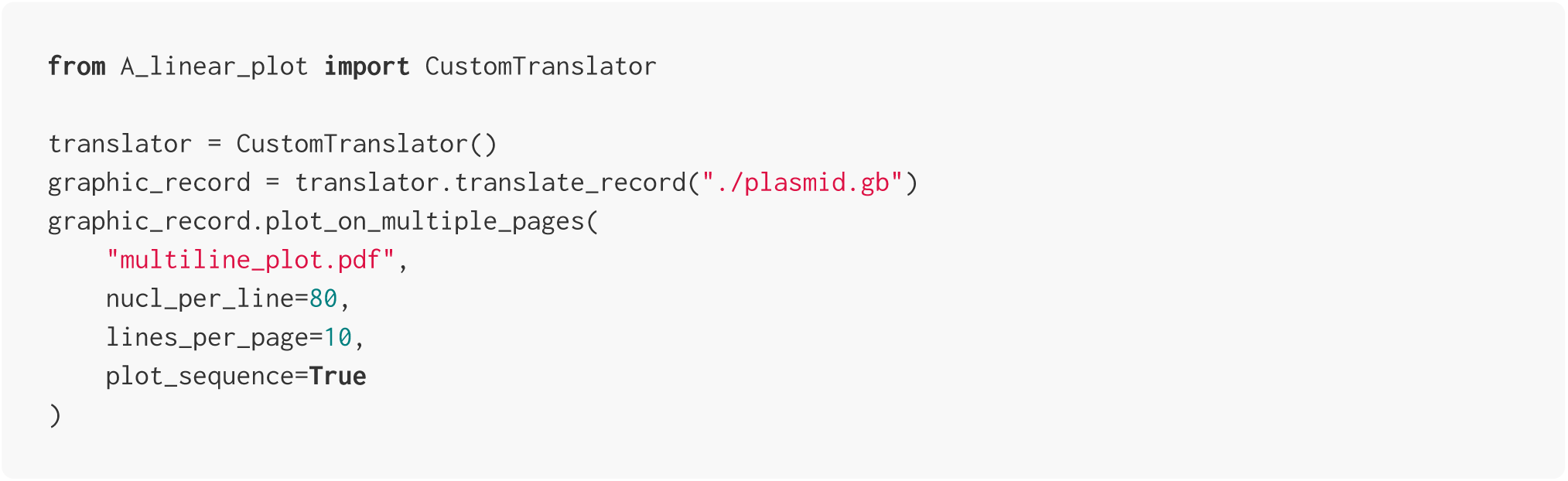

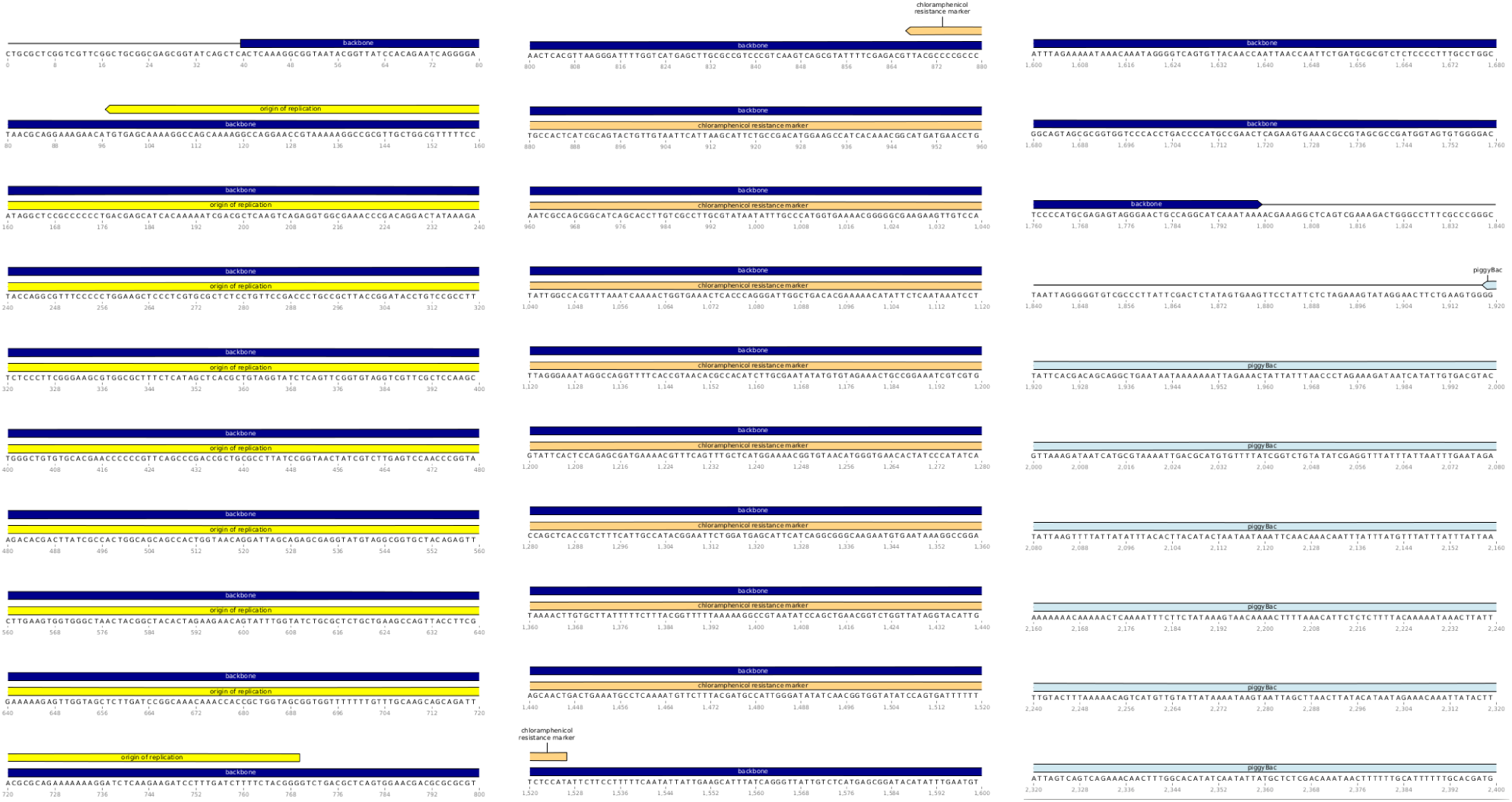

